# DcaP-Family Porins are Required for Carboxylic Acid Utilization and Infection in *Acinetobacter baumannii*

**DOI:** 10.1101/2025.07.02.662759

**Authors:** Hannah R. Noel, Nikita R. Kesav, Jonathan D. Winkelman, Lauren D. Palmer

## Abstract

*Acinetobacter baumannii* is a pathogen of concern and a leading cause of multidrug-resistant healthcare-associated infections. The *A. baumannii* outer membrane is a barrier to antimicrobials and host defenses but must also allow essential nutrients to permeate. Here, we investigate the functional importance of the putative DcaP-family outer membrane porins, which include a proposed vaccine target in *A. baumannii*. All *A. baumannii* genomes surveyed encode multiple DcaP family porins, which we classify in four classes based on protein sequence phylogeny (DcaP1-4). DcaP proteins encoded by species in other genera could not be mapped to these DcaP classes and phylogenetic analysis suggests DcaP1-4 proteins diversified within *Acinetobacter*. Phenotypic array assays and additional experiments show that the DcaP proteins were necessary for growth on multiple di- and tri-carboxylic acids as sole carbon sources, including citric acid and tricarballylic acid. Specifically, DcaP3 was required; however, DcaP1, DcaP2, and DcaP4 were all able to complement the total *dcaP* knockout to varying degrees, suggesting partial functional overlap. Finally, a mutant lacking all DcaP proteins was attenuated in the liver and spleen in a mouse model of bloodstream infection and complemented by expression of DcaP3. However, a Δ*dcaP3* mutant had no defect, demonstrating functional redundancy among DcaP proteins during infection. These findings provide insight on how *A. baumannii* acquires nutrients through the outer membrane barrier and show DcaP proteins are important during infection in specific host niches, validating their potential as a therapeutic target.

## INTRODUCTION

*Acinetobacter baumannii* is a pathogen of critical concern and a major cause of healthcare-associated infections. Isolates of *A. baumannii* are often multidrug resistant (MDR) or extensively drug resistant (XDR) and exhibit resistance to first and last line antibiotics, such as meropenem and colistin, respectively (1). The Centers for Disease Control and the World Health Organization have therefore identified *A. baumannii* as an urgent public health threat, calling for new avenues of treatment (2–4). The bacterial cell envelope is the primary barrier from the environment and protects bacteria from environmental stress including antibiotics and host defenses; however, the cell envelope must also allow acquisition of essential nutrients and cofactors (5,6). The outer membrane (OM) of *A. baumannii* is thought to be less permeable compared to other Gram-negative bacteria such as *E. coli*, contributing to intrinsic antibiotic resistance (7). How small molecule nutrients required for growth transit the impermeable OM in *A. baumannii* is incompletely understood.

The OM of *A. baumannii* is an asymmetric lipid bilayer where the inner leaflet is composed of glycerophospholipids and the outer leaflet of lipooligosaccharides (LOS) (5,8,9). The OM bilayer is occupied by proteins that serve a variety of functions, including nutrient influx. Nutrients typically pass through the OM via active transport by TonB-dependent transporters or passive diffusion through porins, which are transmembrane channels that allow passage of small molecules (6,10). The primary porin in *A. baumannii* is OmpA, which shows slow molecular transport in liposome swelling assays, contributing to the low permeability barrier of the OM (11–13). However, the primary function of OmpA is thought to be structural, providing a physical tether between the OM and the peptidoglycan cell wall in *A. baumannii* and other species (14,15). *A. baumannii* encodes multiple other porins including CarO, an OM porin that selectively allows influx of ornithine and other basic amino acids (16), and the Occ family porins, responsible for the influx of compounds like benzoate, hydroxycinnamate, and other aromatic molecules (17–19). The protein DcaP is a predicted OM porin widely distributed across *Acinetobacter* species named for its localization within a genetic locus for utilization of dicarboxylic acids (*dca*) (20). Notably, a DcaP family protein is among the most abundant OM proteins in *A. baumannii* during rat and mouse infections (21). Due to this fact, multiple groups have studied a DcaP-like protein, which we define below as DcaP3, as a promising vaccine candidate for protection against *A. baumannii* infection (21–25). However, the biological role of DcaP family proteins remains unclear and their role in dicarboxylic acid utilization has not been experimentally interrogated.

First identified in Parke *et al*. as part of the dicarboxylic acid catabolic operon of *Acinetobacter baylyi* ADP1, *dcaP* (putative <u>dic</u>arboxylic <u>a</u>cid <u>p</u>orin) and DcaP-like proteins were thought to be specific to the Moraxellaceae family (20). The structure for one DcaP-like protein has been solved, presenting as a homo-trimeric 16-stranded beta-barrel protein with an extended periplasmic N-terminus that forms a coiled coil (21). Due to the genetic localization and induction of expression with the dicarboxylic acid adipate, *A. baylyi* DcaP porins were proposed to be nonspecific porins that facilitate the uptake of dicarboxylic acids (20); however, a role for DcaP porins in carboxylic acid catabolism has not been reported. Molecular dynamics and electrophysiology studies determined a DcaP-like protein to have an affinity for anionic compounds as substrates, such as the clinically relevant β-lactamase inhibitor sulbactam (21). In other studies, DcaP-like proteins have been implicated in biofilm formation and mucin catabolism (26,27). Here, we describe four classes of DcaP-family porins in *Acinetobacter* and investigate their role in carbon source utilization in *A. baumannii* ATCC 17978 and AB5075 strains.

## RESULTS

### Acinetobacter spp. encode multiple DcaP-family porins

The first DcaP was described in *A. baylyi* as part of a dicarboxylic acid catabolic operon (20), and DcaP-like proteins have been identified in various *A. baumannii* clinical isolates and type-strains (21,23). Proteins designated DcaP-like in *A. baumannii* ATCC 17978 are homologous to *A. baylyi* DcaP, sharing ∼30-35% amino acid identity by Clustal Omega multiple sequence alignment (28). However, given the abundance of DcaP-family porins within individual *Acinetobacter* genomes, we hypothesized that these proteins could be further classified based on protein sequence. Using DcaP and DcaP-like protein sequences from a set of deduplicated *A. baumannii* genomes, four distinct classes of DcaP-family porins were identified that cluster independently, including the original *A. baylyi* DcaP which we display as DcaP1 (Fig. 1A, S1A). We next determined the distribution of DcaP-family porins across the *Acinetobacter* genus. Using a set of 250 deduplicated *Acinetobacter* genomes, 235 of which were *A. baumannii*, DcaP-family porins were shown to be widely conserved across the genus (Fig. S1B). Additionally, *Acinetobacter* species within the *A. calcoaceticus/baumannii* (*acb*) complex/clade of pathogenic *Acinetobacter* appeared to be enriched for having multiple DcaP-family porins compared to non-pathogenic *Acinetobacter* species (Fig. S1B). Among the *A. baumannii* genomes, over 130 of the strains encode four *dcaP* genes (Fig. 1B). In fact, no *A. baumannii* genome analyzed encoded fewer than two *dcaP* genes (Fig. 1B). The type-strain *A. baumannii* ATCC 17978 used for most studies here encodes *dcaP1*, *dcaP2*, and *dcaP3* encoded in three distinct loci (Fig. S2A). To determine if the *dcaP* genes were equally distributed across the species, the number of each DcaP subfamily were tallied for each genome and organized based on how many total DcaP-family porins were present. The four classes of DcaP-family porins were equally likely to be encoded in a given *A. baumannii* genome (Fig. 1B). For example, in most cases where a genome encodes four DcaP-family porins, it was most likely to have one of each variant. Alternatively, in a genome encoding five DcaP-family porins, DcaP2 was most likely to be duplicated. Finally, *A. baumannii* AB5075 encodes *dcaP1-3* and also encodes *dcaP4* in another distinct locus (Fig. S2A). These data suggest that DcaP-family porins are encoded across the *Acinetobacter* genus, and enriched in pathogenic *Acinetobacter,* including *A. baumannii*.

**Figure 1.**
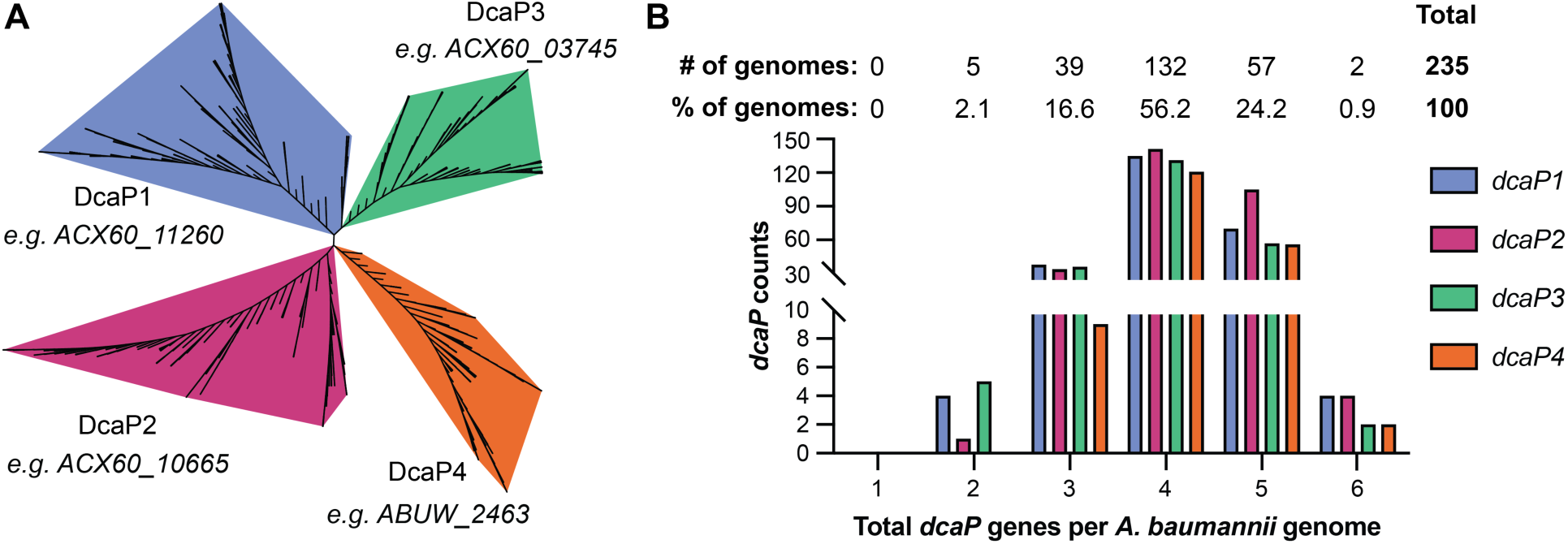
*A. baumannii* DcaP proteins cluster in four classes and are widely distributed among *Acinetobacter*. **(A)** An unrooted tree of DcaP protein sequences from 255 *Acinetobacter* genomes (235 *A. baumannii* genomes). **(B)** Manual counts and distribution of individual DcaP proteins binned by the total number of DcaP proteins in an *A. baumannii* genome.

### DcaP proteins from other bacterial species are distinct from Acinetobacter DcaP proteins

To investigate the evolutionary history of DcaP-family porins in *Acinetobacter* and determine if DcaP diversified within *Acinetobacter* or across taxa, orthologues in other bacterial taxa were identified through KEGG. Bacterial taxa were selected for further analysis if the DcaP-like protein was encoded in more than half of the genomes for a given genus (29,30). A rooted phylogenomic tree was generated to demonstrate the evolutionary distance between *Acinetobacter* and the bacteria containing a DcaP orthologue (Fig. 2A). Orthologues of DcaP-family porins were found in distantly related organisms, such as *Shewanella spp.* and *Xanthomonas spp.,* albeit generally in fewer numbers (Fig. 2A). These DcaP orthologues could not be further classified into one of the four DcaP classes defined in *A. baumannii*. This suggests that the four DcaP classes of proteins defined here are unique to *Acinetobacter spp*. To better understand the relationship between *Acinetobacter* DcaP-family porins and proteins designated DcaP-like from distantly related species, an unrooted tree of protein sequences was generated. This analysis revealed that DcaP-family porins from AB5075, the included reference *A. baumannii* strain which encodes DcaP1-4, were more similar to each other than to non-*Acinetobacter* DcaP-family porins (Fig. 2B). This suggests that diversification of the four DcaP classes defined here occurred within *Acinetobacter*. In contrast to *A. baumannii*, multiple DcaP-like proteins encoded in other organisms were sequence divergent, such as in *Pseudoxanthomonas daejeonensis*, *Stenotrophomonas spp*., *Shewanella psychrotolerans*, and *Marinobacter spp.* (Fig 2B). Comparison of the species phylogeny and DcaP protein phylogeny suggest there may have been occurrences of horizontal gene transfer in other species outside *Acinetobacter*. Overall, these data suggest that DcaP-like proteins are encoded by species across Pseudomonodota and that DcaP diversification has occurred within the *Acinetobacter* clade.

**Figure 2.**
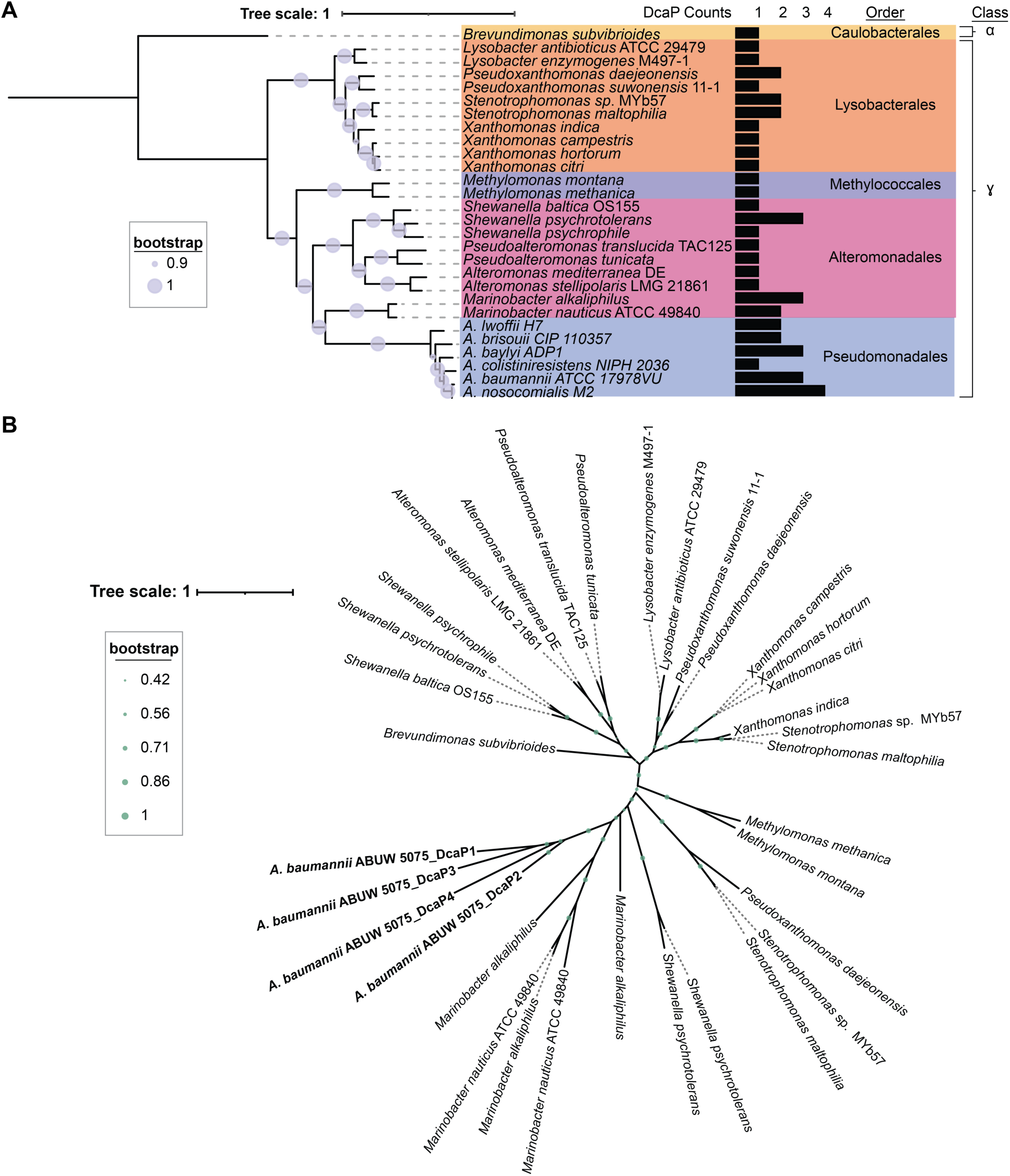
*A. baumannii* DcaP-family proteins cluster together compared to non-*Acinetobacter* DcaP proteins. **(A)** A rooted tree of DcaP containing species outside of *Acinetobacter* and some *Acinetobacter spp.* depicting class, order, and number of encoded DcaP proteins. **(B)** An unrooted tree of DcaP protein sequences from non-*Acinetobacter* species with AB5075 as the *A. baumannii* reference. Tree scale in amino acid substitutions.

### DcaP family porins are necessary for growth on select di- and tri-carboxylic acids

Next, we hypothesized that the DcaP-family porins are involved in small molecule uptake. One study suggested that DcaP3 may translocate the clinically relevant beta-lactamase inhibitor sulbactam based on electrophysiology and applied field simulation data (21). Many *A. baumannii* isolates are intrinsically resistant to the beta-lactam ampicillin due to the expression of beta-lactamases and sulbactam has been shown to resensitize resistant isolates to beta-lactam antibiotics (31,32). We reasoned that if DcaP-family porins facilitated sulbactam entry, a mutant lacking all *dcaP* genes would exclude sulbactam from entering the cell and result in increased resistance to the beta-lactam ampicillin while a wildtype strain would exhibit re-sensitization to ampicillin. We therefore constructed a triple mutant in ATCC 17978 lacking all encoded DcaP proteins, Δ*dcaP1*Δ*dcaP2*Δ*dcaP3* (ΔΔΔ*dcaP1-3*). However, the wildtype and the ΔΔΔ*dcaP1-3* mutant strains exhibited similar susceptibility to ampicillin/sulbactam treatment (Fig. S2). These data suggest that DcaP-family porins are not required for sulbactam activity.

We next hypothesized that DcaP-family porins may be important for growth on certain nutrient sources such as dicarboxylic acids as previously suggested (20). The growth of wildtype and ΔΔΔ*dcaP1-3* strains were compared using the high-throughput Biolog PM system to screen for carbon source utilization. Multiple compounds were identified on which ΔΔΔ*dcaP1-3* grew more than wildtype and compounds on which wildtype was able to grow but ΔΔΔ*dcaP1-3* had a defect (Fig. 3A-B). The screen suggested ΔΔΔ*dcaP1-3* had increased growth compared to wildtype on amino acids such as glutamic acid and glutaminic acid (Fig. 3A-B); however, validation showed that there was no difference in growth between wildtype and ΔΔΔ*dcaP1-3* (Fig. S3B). Glutamic acid was thus used as a control for subsequent assays to ensure growth of strains in minimal media. Butyric acid also appeared to better support growth of ΔΔΔ*dcaP1-3* and was analyzed in subsequent assays. Many of the compounds on which WT grew better than ΔΔΔ*dcaP1-3* were carboxylic acids, including simple carboxylic acids, hydroxy carboxylic acids, and sugar acids (Fig. 3A-B). These data suggest that DcaP-family porins may allow the influx of carboxylic acid-containing compounds, as previously predicted (20).

**Figure 3.**
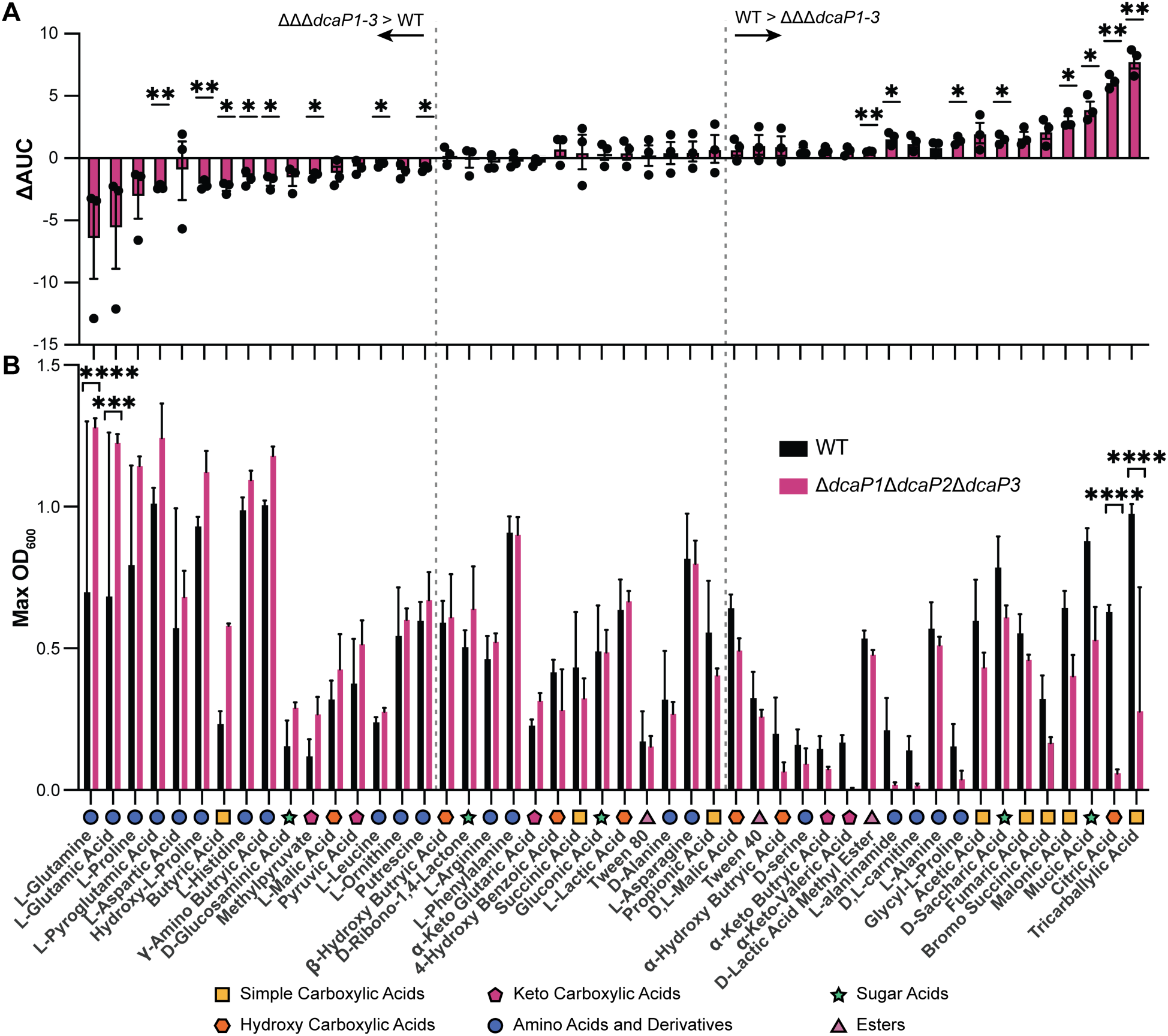
Wildtype *A. baumannii* grows better than ΔΔΔ*dcaP1-3* mutant on certain di- and tricarboxylic acids as sole carbon sources. Biolog plates PM1 and PM2a were resuspended in M9 minimal media without a carbon source and inoculated with either WT or the **ΔΔΔ***dcaP1-3* mutant. (**A**) Difference in area under the curve (**Δ**AUC) was calculated for WT – **ΔΔΔ***dcaP1-3* mutant. Each point is from one experiment with 1 biological replicate per strain. Significance is by one sample t-test compared to 0. (**B**) Maximum optical density at 600 nm (OD_600_) is shown. Significance is by two-way ANOVA with Sidak’s multiple comparisons. Experiments were conducted three times with an n=1 for a total of n=3. Data are mean +/- SD. \**P* < 0.05, \*\**P* < 0.01, ***P < 0.001, ****P < 0.0001

### DcaP3 is important for growth on multiple carboxylic acid carbon sources

To verify the Biolog results and further determine functional importance of the DcaP classes, growth curves using ΔΔΔ*dcaP1-3,* Δ*dcaP1*Δ*dcaP2*, Δ*dcaP1*Δ*dcaP3*, and Δ*dcaP2*Δ*dcaP3* and carboxylic acids as sole carbon sources were performed. The strains were also assayed on glutamic acid and succinic acid as controls for growth in minimal media, and butyric acid as a potential carbon source in which ΔΔΔ*dcaP1-3* grew to higher yield than WT. There were no differences in growth in glutamic acid, and potential minor defects for growth of the ΔΔΔ*dcaP1-3* strain in succinic acid and butyric acid. This showed that the screen did not identify any conditions where the ΔΔΔ*dcaP1-3* strain grew more than WT. By contrast, mutants lacking DcaP3 were defective for growth on citric acid, tricarballylic acid, and had a minor defect on mucic acid compared to the WT (Fig. 4A). We were unable to identify conditions in which DcaP1 or DcaP2 were required for growth. Similarly, a single Δ*dcaP3* mutation conferred a defect in growth on citric acid, tricarballylic acid, and mucic acid, whereas single mutants of Δ*dcaP1* or Δ*dcaP2* showed no defect (Fig. 4B).

**Figure 4.**
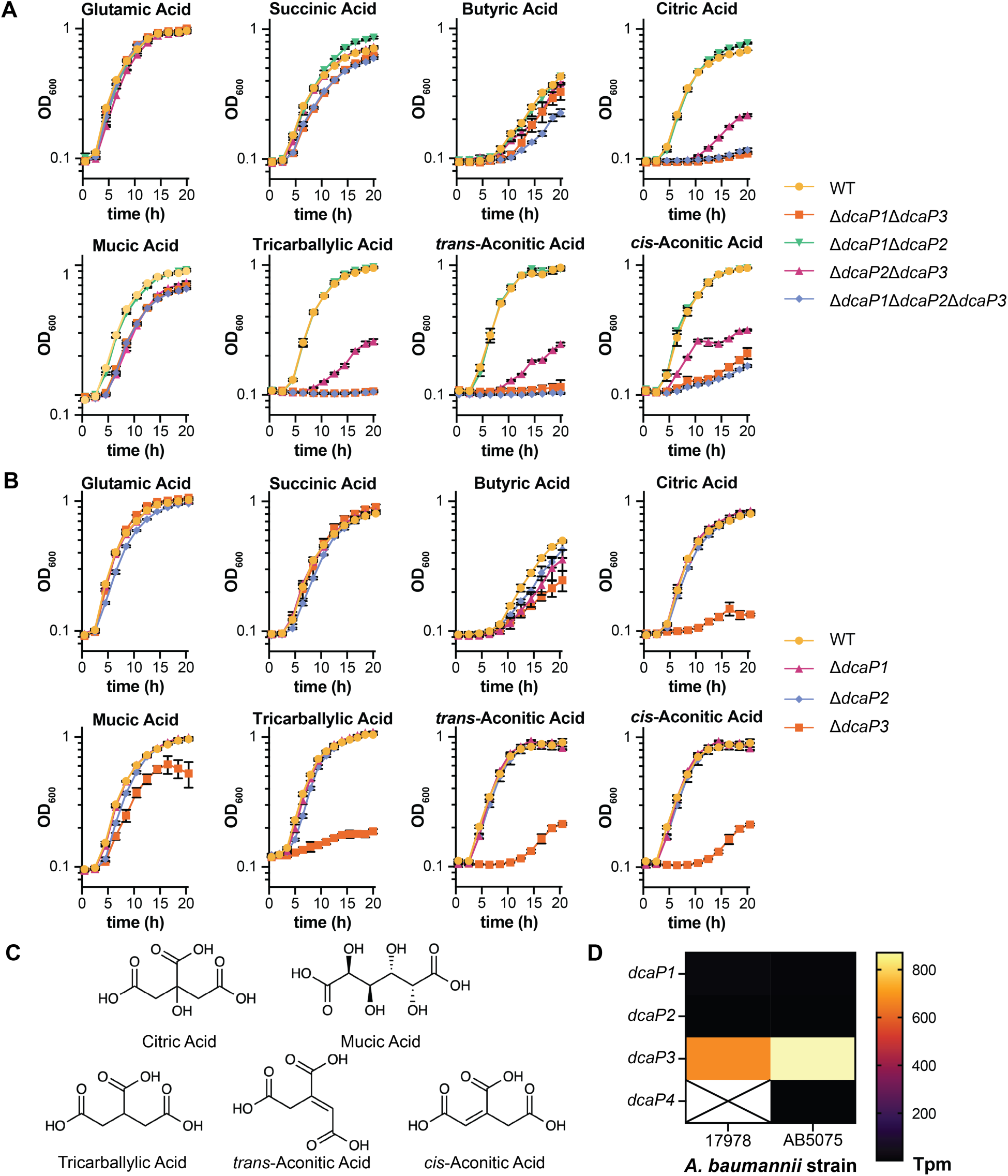
DcaP3 is important for growth on the carboxylic acid carbon sources citric acid, tricarballylic acid, mucic acid, and *cis*- and *trans*-aconitic acid. **(A)** Wildtype, double, and the triple ΔΔΔ*dcaP1-3* mutants were grown in M9 media with the indicated compound as the sole carbon source. Growth was monitored by optical density at 600 nm (OD_600_). **(B)** Wildtype, the ΔΔΔ*dcaP1-3* mutant, and the ΔΔΔ*dcaP1-3* mutant complemented with *dcaP3* were grown on the indicated compound as the sole carbon source. **(C)** The chemical structures of the relevant carbon source compounds. **(D)** Expression of the *dcaP* genes in *A. baumannii* 17978 and AB5075 from (34). Average transcripts per million (Tpm) reads. (A-B) Data are mean +/- SEM and n=2-3. Experiments were repeated at least twice with similar results.

A recent study in *A. baylyi* found that tricarballylic acid metabolism is closely linked with the metabolism of *cis*- and *trans*-aconitic acid (33). The genetic organization of the tricarballylic acid catabolic genes is conserved in *A. baumannii* compared to *A. baylyi* (Fig. S4A). Since the data thus far suggest DcaP3 is important for growth on tricarballylic acid, we hypothesized that DcaP3 may also be important for *cis*- and *trans*-aconitic acid growth. Mutants lacking DcaP3 had a defect for growth on both *cis*- and *trans*-aconitic acid (Fig. 4A) and the Δ*dcaP3* single mutant had a defect in growth on both *cis*- and *trans*-aconitic acid compared to wildtype (Fig. 4B). To highlight the similarities in the carbon sources in which DcaP3 is important for growth, the chemical structures for each carbon source are outlined in Fig. 4C. Given the lack of identifiable phenotype for Δ*dcaP1* or Δ*dcaP2*, transcript reads from previously published datasets were analyzed to determine if *dcaP1* or *dcaP2* are expressed (34). In *A. baumannii* 17978, *dcaP3* was expressed at a high level in untreated lysogeny broth while *dcaP1* and *dcaP2* were not expressed; in *A. baumannii* AB5075, *dcaP3* was similarly highly expressed while *dcaP1*, *dcaP2*, and *dcaP4* were not expressed (Fig. 4D) (34). Similar results were observed in another *A. baumannii* 17978 dataset (35). Taken together, these data suggest that DcaP3 is highly expressed and important for growth using multiple di- and tri-carboxylic acids as sole carbon sources.

### DcaP3 is required for growth on di- and tri-carboxylic acids in a recent clinical isolate

To test if DcaP3 is important for growth on di- and tri-carboxylic acids in a more recent clinical isolate, a Δ*dcaP3* mutant was generated in *A. baumannii* AB5075, a member of the widely circulating clonal complex 1 that was isolated in 2008 and also serves as a commonly used reference strain (36). An *A. baumannii* AB5075 mutant lacking *dcaP3* was defective for growth on citric acid, tricarballylic acid, mucic acid, and *cis*- and *trans*-aconitic acid compared to wildtype (Fig. 5). This suggests that the requirement for DcaP3 for growth on select carboxylic acids as sole carbon sources *in vitro* is conserved among *A. baumannii* isolates.

**Figure 5.**
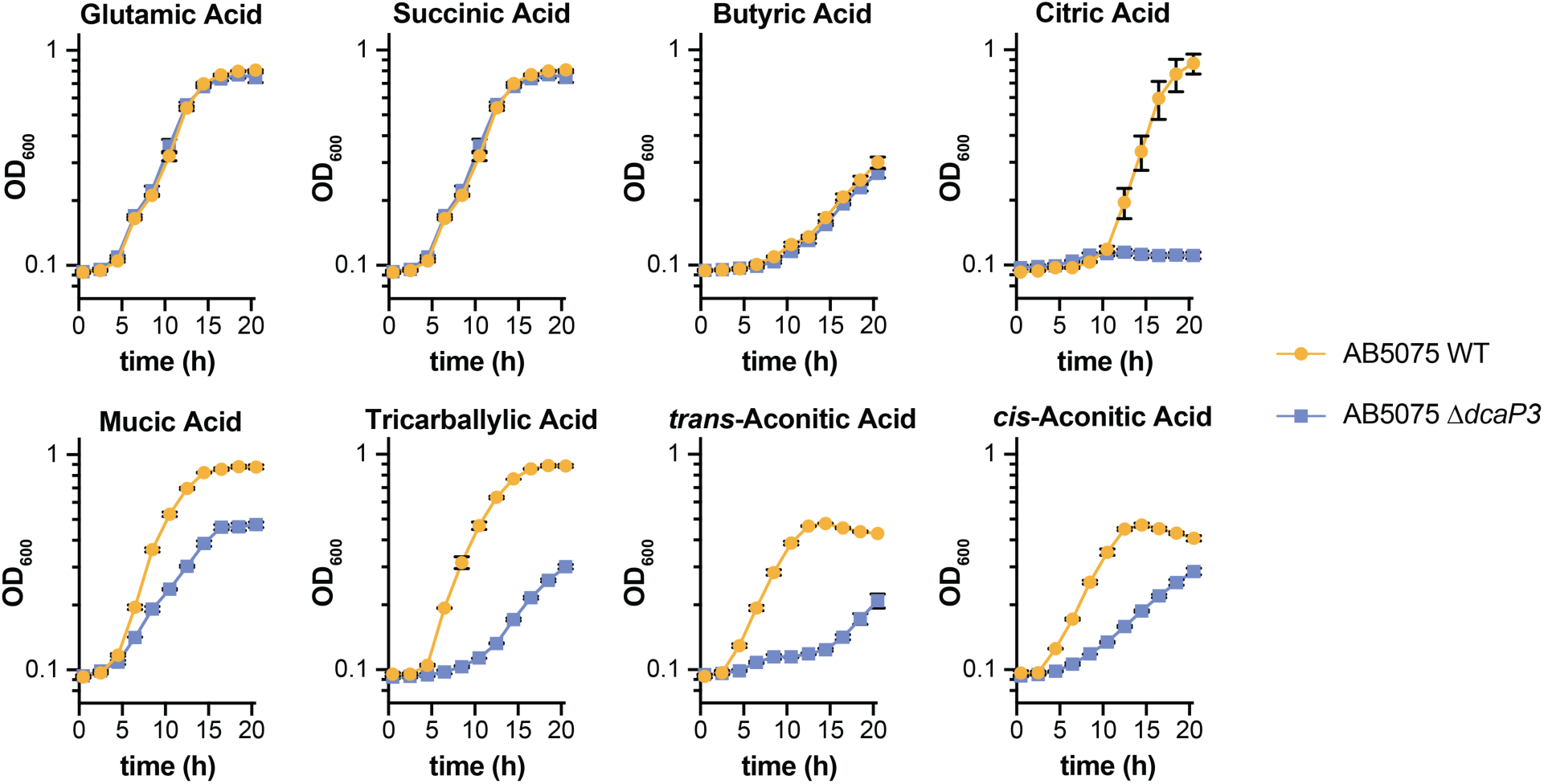
*A. baumannii* AB5075 Δ*dcaP3* is defective for growth on citric acid, tricarballylic acid, mucic acid, and *cis*- and *trans*-aconitic acid. ***A.*** *baumannii* AB5075 wildtype (WT) and Δ*dcaP3* mutant strains were grown in M9 media with the indicated compounds as the sole carbon source. Growth was monitored by optical density at 600 nm (OD_600_). Data are mean +/-SEM and n=3. Experiments were repeated twice with similar results.

### DcaP1-4 can complement ΔΔΔdcaP for growth on some carboxylic acids

Thus far, genetic disruption in *A. baumannii* ATCC 17978 and AB5075 suggested DcaP3 was the primary DcaP important for utilization of certain di- and tri-carboxylic acids. However, published expression analysis suggests only DcaP3 is expressed in laboratory conditions (Fig. 4D) (34,35). We hypothesized that individual DcaP proteins may complement the *A. baumannii* 17978 ΔΔΔ*dcaP1-3* growth defects if expressed from the *dcaP3* promoter. Therefore, chromosomal mini-Tn*7* (mTn*7*) complementation constructs were constructed expressing *A. baumannii* 17978 *dcaP1*, *dcaP2*, or *dcaP3* or *A. baumannii* AB5075 *dcaP4* from the *A. baumannii* 17978 *dcaP3* promoter. Consistent with previous observations, the ΔΔΔ*dcaP* mutant was unable to grow using citric acid, tricarballylic acid, or *cis*- and *trans*-aconitic acid and had a slight defect on succinic acid, butyric acid, and mucic acid (Fig. 6). Growth was restored in succinic acid and butyric acid by expressing *dcaP4* (Fig. 6), suggesting DcaP4 can contribute to succinic acid and butyric acid transport. Similarly, growth on citric acid, mucic acid, tricarballylic acid, and trans- and cis-aconitic acid was restored to the ΔΔΔ*dcaP1-3* mutant by expressing *dcaP2, dcaP3*, or *dcaP4* (Fig. 6). This suggests that DcaP2, DcaP3, and DcaP4 have overlapping functions in at least some *A. baumannii* strains and conditions. Expression of *dcaP1* in the ΔΔΔ*dcaP1-3* strain did not restore growth in succinic acid, butyric acid, citric acid, mucic acid, or tricarballylic acid but did restore growth in *trans*- and *cis-*aconitic acid, suggesting a largely non-overlapping functional role for DcaP1.

**Figure 6.**
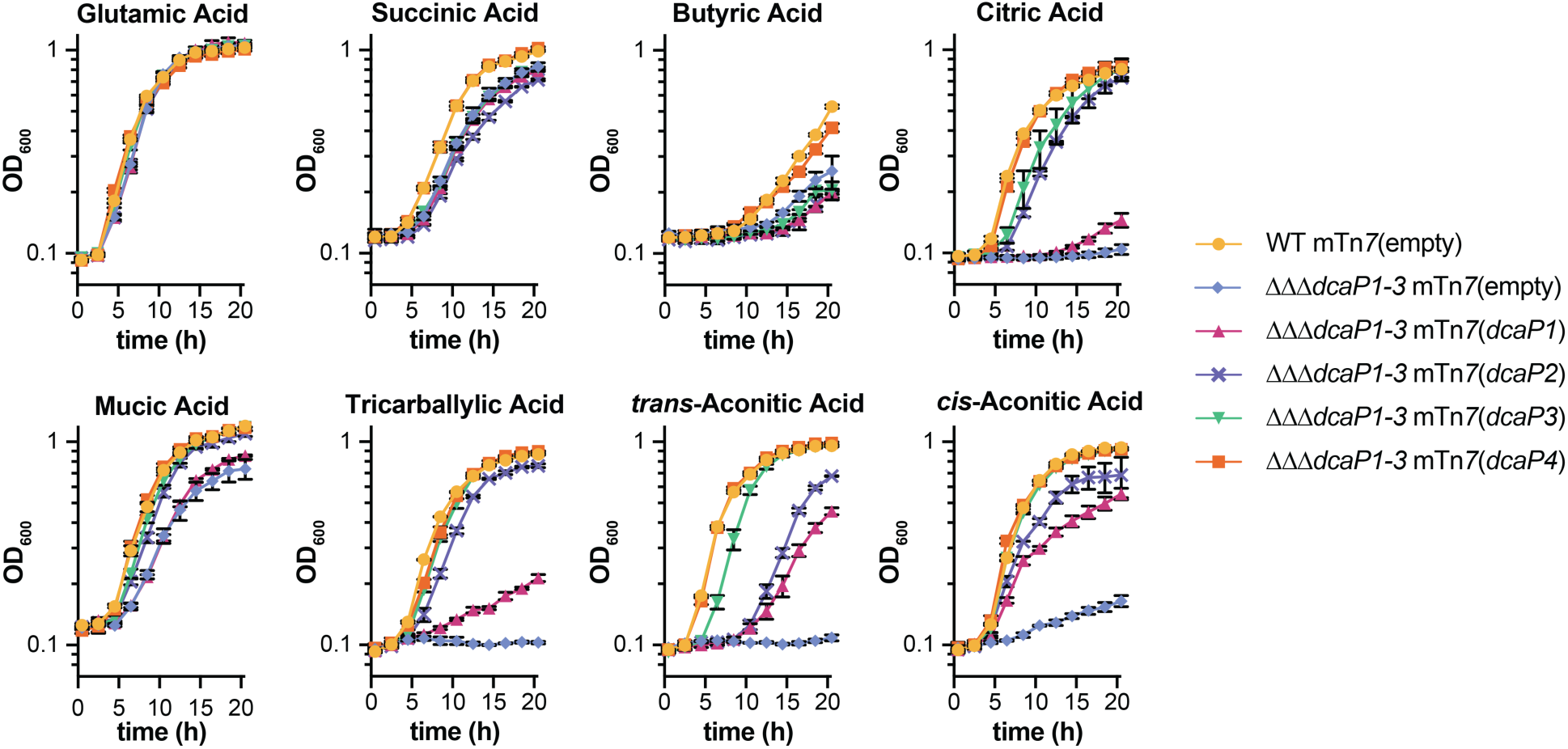
DcaP1, DcaP2, DcaP3, and DcaP4 can complement growth of *A. baumannii* ATCC 17978 ΔΔΔ*dcaP1-3* with certain sole carbon sources. Wildtype, the ΔΔΔ*dcaP1-3* mutant, and the ΔΔΔ*dcaP1-3* mutant complemented with *dcaP1, dcaP2, dcaP3,* or *A. baumannii* AB5075 *dcaP4* were grown in M9 media with the indicated compound as the sole carbon source. Growth was monitored by optical density at 600 nm (OD_600_). Data are mean +/- SEM and n=3. Experiments were repeated at least twice with similar results.

### DcaP3 is sufficient but not required for virulence in the liver and spleen during bloodstream infection of mice

Carbon source utilization can be a critical determinant of pathogenesis for *A. baumannii* and other bacterial species in different host niches and DcaP3 has been proposed as an *A. baumannii* vaccine candidate (21–25,37,38). Therefore, we sought to determine if DcaP-family porins are important for *A. baumannii* pathogenesis in a murine model of bloodstream infection. There was no defect of the ΔΔΔ*dcaP1-3* mutant compared to wildtype in lung, kidney, or heart (Fig. 7A). By contrast, there was a defect for the ΔΔΔ*dcaP1-3* mutant compared to wildtype in the spleen and liver (Fig. 7A). The defect in the spleen and liver was reversed by complementing *dcaP3* in the chromosome of the ΔΔΔ*dcaP1-3* mutant, showing DcaP3 is sufficient for proliferation in the liver and spleen (Fig. 7B). However, a single Δ*dcaP3* mutant had no defect during infection (Fig. 7C). This suggests that in the host, DcaP1 and/or DcaP2 are expressed and able to functionally complement the lack of DcaP3 in *A. baumannii* 17978. Together these data show that DcaP-family porins are important in specific host niches during infection.

**Figure 7.**
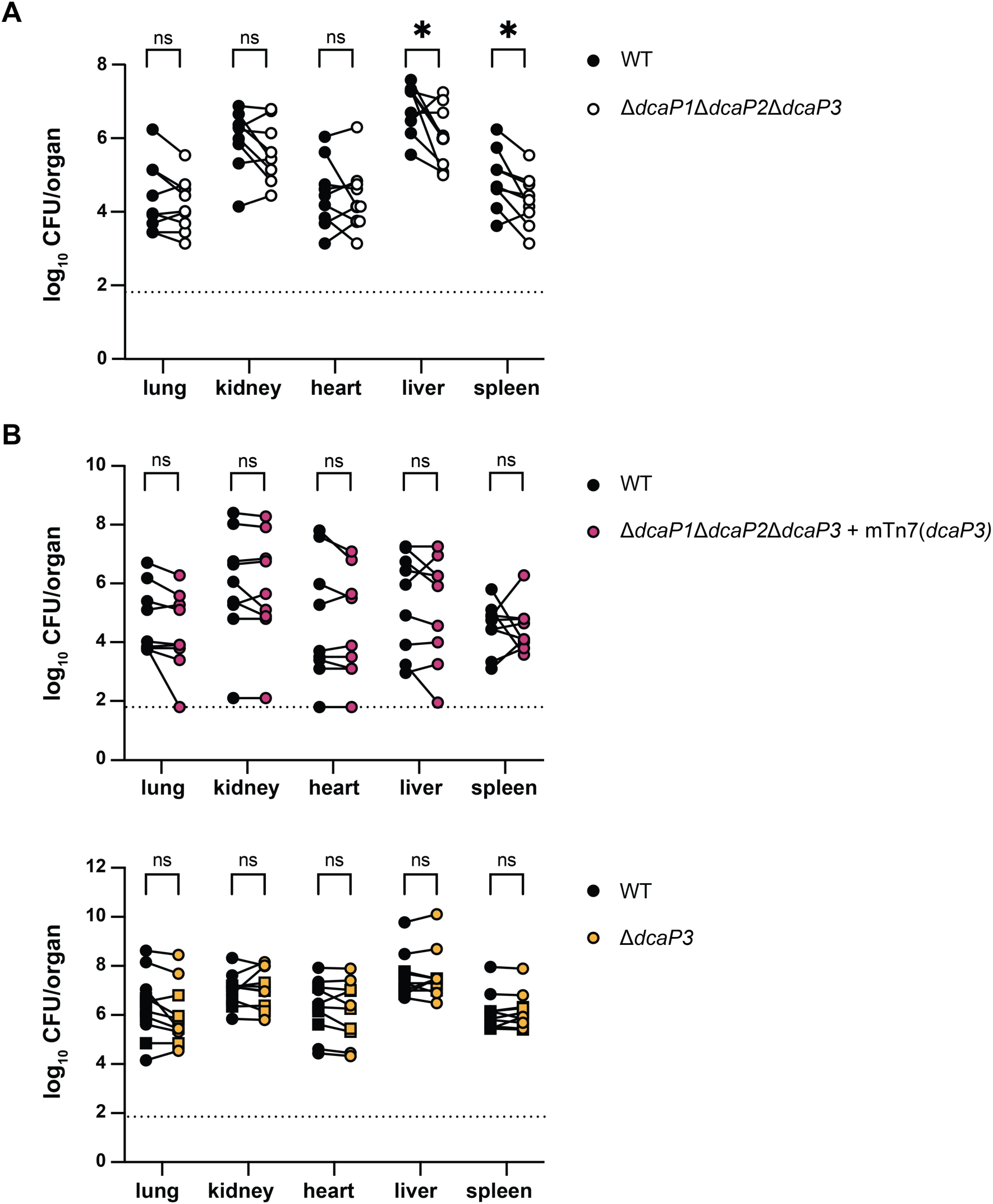
DcaP3 is important for infecting the liver and spleen in a model of bloodstream infection. **(A-C)** Wildtype (WT) and the indicated strains were retro-orbitally inoculated into Swiss Webster mice in a 1:1 ratio. At 24 h post-infection, organs were harvested and bacterial CFU were enumerated. The dotted lines indicate the limit of detection. Each point represents one strain enumerated from one mouse; n=9-10. Circle symbols are female mice and square symbols are male mice. (A-B) Data are combined from two experiments. (C) Data are from one experiment. Statistics are Wilcoxon non-parametric test. \**P* < 0.05.

## DISCUSSION

The OM protects *A. baumannii* from antimicrobials and other host stresses but must permit influx of nutrients essential for replication. How nutrients pass through the *A. baumannii* OM is not well understood. Here, we designate classes of a family of predicted porins, DcaP1-4, that are widely distributed across *Acinetobacter spp.* and show DcaP-family porins are required for growth on multiple carboxylic acids.

DcaP-family porins have been identified in genetic screens for biofilm formation, desiccation, and mucin catabolism, largely referred to as DcaP-like proteins without further distinction (26,27,39–41). We show that *A. baumannii* DcaP-family porins can be grouped into four classes based on protein sequence (Fig. 1A, S1A). DcaP-encoding genes are broadly conserved among *Acinetobacter* spp, and particularly in *A. baumannii* and the *acb* complex (Fig. 1B, Fig. S1B). While DcaP-family porins can be found in distantly related organisms (Fig. 2A), the DcaP1-4 classes we define in *A. baumannii* cluster together (Fig. 2B). These findings suggest DcaP1-4 diversified within the *Acinetobacter* clade.

Antibiotic resistance in *A. baumannii* is a critical public health problem due to the emergence of XDR isolates. Various porins in *A. baumannii* have been implicated in the uptake of toxic molecules including antibiotics (12,42–44). DcaP3 was proposed to allow entry for the beta-lactamase inhibitor sulbactam (21). Here, we show that a ΔΔΔ*dcaP1-3* mutant displays similar susceptibility to ampicillin/sulbactam as the wildtype *A. baumannii* 17978, suggesting that DcaP-family porins are not required for sulbactam function (Fig. S2). However, additional studies would be required to determine whether sulbactam may enter via DcaP-family porins in addition to other outer membrane porins.

The transport of essential nutrients across the cell envelope is incompletely understood in *A. baumannii*. In general, *A. baumannii* utilizes organic acids and amino acids as carbon sources and does not utilize sugars such as glucose (37,45). Accordingly, multiple OM channels have been identified that are important for growth on specific carbon sources including ornithine, arginine, glutamic acid, and benzoic acid (16–19,46). One study in ATCC 19606 found that *dcaP1* (*A1S_1380*) was upregulated in the presence of mucin (26), suggesting that the host environment may affect *dcaP* gene regulation. In a Biolog screen, we identified multiple carboxylic acid carbon sources that supported significantly more growth of the wildtype compared to the ΔΔΔ*dcaP1-3* mutant (Fig. 3), suggesting DcaP-family porins are important for their diffusion through the OM.

The data here suggest DcaP3 is responsible for the influx of carboxylic acid carbon sources citric acid, mucic acid, tricarballylic acid, and *cis*- and *trans*-aconitic acid in *A. baumannii* ATCC 17978 and AB5075 in the *in vitro* conditions tested, as mutants lacking DcaP3 show decreased growth on these substrates (Fig. 4-5). Phylogenomics analysis found three *A. baumannii* strains lacking *dcaP3*: *A. baumannii* E-011922, *A. baumannii* VB82, and *A. baumannii* RBH2 (Fig. S1B) (47–49). Of these, carbon utilization data were reported for *A. baumannii* E-011922, an environmental isolate that retains the ability to grow on citric acid and mucic acid despite not encoding DcaP3 (48). Similarly, *A. baylyi* can grow on citric acid and tricarballylic acid and does not encode DcaP3 but rather DcaP1, DcaP2, and DcaP4 (33). This suggests that in strains that lack DcaP3, other DcaP-family porins are likely expressed at higher abundance and allow transport of these carboxylic acids.

Citric acid is an intermediate of the tricarboxylic acid (TCA) cycle, a ubiquitous pathway for energy generation. Studies quantifying citric acid in rat and mouse tissues report it is 0.15-0.3 µmol/g in the liver and 0.27-0.45 µmol/g in the kidney (50–52), suggesting citric acid is abundant in some tissues. Despite this, many bacteria cannot utilize citric acid as a carbon source and how citric acid permeates the OM is unknown (53). Mutants in *P. aeruginosa* OpdH were defective for growth on *cis*-aconitic acid, but not on citric acid, suggesting they encode other citric acid transporters (54,55). A ΔΔΔ*dcaP1-3* mutant displayed a strong defect for growth on citric acid that can be rescued by complementing with *dcaP2*, *dcaP3* or *dcaP4* (Fig. 4A, 4B, Fig. 5). Therefore, encoding multiple porins for OM citric acid passage may be conserved across bacterial genera.

Mucic acid, or galactaric acid, is a sugar acid derived from the oxidation of galactose and is unrelated to mucin. Mucic acid is not typically found in the host; but, some antibiotic treatments result in increased mucic acid in the human gut (56–58). However, only a subset of *A. baumannii* strains, including ATCC 17978, encode the D-galactaric acid dehydrogenase required for growth on mucic acid (59). The digestive tract is a reservoir for *A. baumannii* colonization that increases the potential for infection and transmission and *A. baumannii* ATCC 17978 can asymptomatically colonize the mouse gut (60–65). Thus, mucic acid may be used as carbon source by some *A. baumannii* in the inflamed gut. Our data suggest that DcaP2, DcaP3 or DcaP4 are important for growth on mucic acid.

The catabolism of *cis*- and *trans*-aconitic acid has been described in *A. baylyi*, and the catabolism of tricarballylic acid in *A. baylyi* and *Salmonella* (33,66,67). However, the OM porin required for growth on these carbon sources in *Acinetobacter* was unknown. Our data suggest that DcaP2, DcaP3, and DcaP4 porins are important for growth on tricarballylic acid, and that DcaP1 can also support growth on *cis*- and *trans*-aconitic acid.

Overall, our data suggests that DcaP3 is the major porin for passive transport of specific carboxylic acid carbon sources in the *in vitro* conditions tested. The requirement for DcaP3 in growth on select di- and tri-carboxylic acids is conserved in the more recent clinical isolate *A. baumannii* AB5075, as a Δ*dcaP3* mutant had reduced growth in these conditions (Fig. 6). Published RNA sequencing data show that *dcaP3* is the only *dcaP* gene expressed under laboratory conditions tested, consistent with the defects for the Δ*dcaP3* single mutants in *A. baumannii* 17978 and AB5075 strain backgrounds (Fig. 4B, Fig. 5) (34,35).

During infection, bacteria must scavenge nutrients from the host to replicate. The *A. baumannii* ATCC 17978 ΔΔΔ*dcaP1-3* mutant had a defect compared to wildtype in the mouse liver and spleen that was complemented by expression of *dcaP3* (Fig. 7A-B). However, the Δ*dcaP3* single mutant had no defect (Fig. 7C). Therefore, DcaP3 passive import of carboxylic acid carbon sources is sufficient but not necessary for pathogenesis in the liver and spleen. This suggests the host environment may lead to increased expression of other *dcaP* genes; regulation of *dcaP* expression and the molecular basis for substrate specificity is an area for future study. Additionally, future study will be required to dissect the relevant carboxylic acid carbon sources and DcaP substrates important for *A. baumannii* infection. There is little known about the role for these carboxylic acid carbon sources during bacterial infection. Citric acid has been shown to serve as a signaling molecule for *Staphylococcus aureus* pathogenesis; citric acid was increased in the lung at 4 weeks during *Mycobacterium tuberculosis* infection, and citric acid utilization promoted *Vibrio cholerae* small intestine colonization (68,69). The *Klebsiella pneumoniae* citric acid synthase GltA has been shown to be important due to amino acid biosynthesis required in the spleen, liver, and gut, but dispensable in the bloodstream (70). Succinic acid has also been shown to be an important carbon source for *Pseudomonas aeruginosa* during infection (71). Alternatively, DcaP-family porins may play another role in pathogenesis. For example, porins in *A. baumannii* have demonstrated roles for the adhesion or invasion of eukaryotic cells (13,72–75). DcaP3 has been proposed as a vaccine candidate due to its high expression in infection models and immunogenicity (21–25). These data suggest that vaccine strategies to functionally block DcaP proteins must block multiple DcaP proteins to affect *A. baumannii* virulence.

In summary, we further classify a family of porins previously termed DcaP-like into four classes in *Acinetobacter* spp. This classification allows for the differentiation between DcaP-like proteins and the potential to determine individual functions for DcaP1-4 in *Acinetobacter* spp. DcaP3 was critically important for growth on citric acid, tricarballylic acid, mucic acid, and *cis*-and *trans*-aconitic acid as sole carbon sources. In a murine model of bloodstream infection, DcaP3 was sufficient but not necessary for infection, suggesting functional overlap among DcaP porins in the host. Expressing *dcaP2*, *dcaP3*, or *dcaP4* complemented growth of the *A. baumannii* 17978 ΔΔΔ*dcaP1-3* mutant on all carbon sources tested and expression of *dcaP1* complemented growth on *trans*- and *cis*-aconitic acid (Fig. 5). Our data therefore provides a deeper understanding of how *A. baumannii* acquires nutrients including in the host.

## MATERIALS AND METHODS

### Bacterial strain construction and growth

All bacterial strains and plasmids used in this study are listed in Tables S1 and S2, respectively. The wildtype strain was *A. baumannii* ATCC 17978VU or AB5075 (36,76,77). Oligonucleotides are listed in Table S3 and were purchased from Integrated DNA Technologies (Coralville, IA). Strains were grown in lysogeny broth (LB; Miller) or on LB with 1.5% (wt/vol) agar unless indicated otherwise. Antibiotics were used at the following concentrations: carbenicillin (Carb), 75 mg/L; kanamycin (Kan), 40 mg/L; chloramphenicol, 15 mg/L; hygromycin (Hyg) 300 mg/L. For deletion constructs, 1000 bp of 3’ and 5’ flanking DNA to the gene of interest was amplified using Q5 High Fidelity 2x Master Mix [New England Biolabs (NEB), Ipswich, MA] from *A. baumannii* ATCC 17978VU or AB5075. The vector pFLP2 was digested with BamHI and KpnI restriction enzymes (NEB, Ipswitch, MA) and gel purified (78). The kanamycin or hygromycin cassette was amplified from pKD4 or pLDP357, respectively (79). A hygromycin-resistant excisable cassette was generated by replacing kanamycin resistance in pKD4 with hygromycin resistance amplified from an *A. baumannii* AB5075 transposon T101 mutant (36,77) with XR185/XR186 oligonucleotides and ligation with NEBuilder^®^ HiFi DNA Assembly Master Mix (NEB, Ipswich, MA) to generate pLDP357. Fragments were ligated together using HiFi ligation mix (NEB, Ipswitch, MA), transformed into chemically competent *E. coli* DH5α, and plated to selective media. Transformants were confirmed to contain the insert by PCR. To generate the knockout strains, a triparental conjugation using helper *E. coli* strain HB101 containing pRK2013 was performed. Single-crossover merodiploids containing the integrated pFLP2 vector were chosen based on a Kan^R^ Carb^R^ or Hyg^R^ and sucrose-sensitive (10% w/v; Suc^S^) phenotype. Putative merodiploids displaying were verified by PCR. Merodiploids were then grown on LB agar, resuspended, and plated to LB agar or LB agar with 10% sucrose to select for colonies with the second crossover event. Colonies were then screened for a Kan^R^ Carb^S^ Suc^R^ or Hyg^R^ Suc^S^ phenotype. Mutants were confirmed via PCR.

For chromosomal integration constructs, DNA from ATCC 17978VU or AB5075 was amplified using Q5 High Fidelity 2x Master Mix (NEB, Ipswich, MA). The mTn*7* pKNOCK vector was digested with BamHI and KpnI prior to ligation using HiFi ligation mix (NEB, Ipswitch, MA) and transformation into chemically competent *E. coli* DH5α λ*pir116*^+^ (80). To generate chromosomally integrated mTn7 constructs, a modified four-way parental mating was performed using helper *E. coli* strain HB101 containing pRK2013 and *E. coli* DH5α containing pTNS2 (81). Colonies were screened by PCR for integration into the correct chromosomal locus at the *att* site. All plasmid constructs were verified by whole plasmid sequencing before transforming into *A. baumannii* (Plasmidsaurus, South San Francisco, CA).

The *A. baumannii* AB5075 wildtype and Δ*dcaP3*::hyg mutant were verified to have the virulent/opaque (VIR-O) phenotype on 1/2X LB agar plates (82,83).

### Phylogenetic tree construction

To identify DcaP orthologs and classify them into hierarchical orthogroups (HOGs), OrthoFinder v2.5.5 was used (84). Two datasets were analyzed: (i) a 253-strain dataset including 235 *A. baumannii* strains deduplicated as previously described (85); (ii) a 28-species dataset that included multiple *Acinetobacter* and a broader range of gammaproteobacteria rooted by one alphaproteobacterial. The protein accession numbers and genome accession numbers are listed in Table S4.

Non-Acinetobacter species were selected based on the conservation of DcaP orthologs across the genus or species indicated by KEGG. Specific organisms were selected to portray a representative distribution of DcaP orthologs across bacteria, a group of 22 species. The FastTree v2.1.11-derived (86), manually rooted species tree was used as an input for OrthoFinder HOG inference (84,87), ensuring accurate classification of gene duplication and speciation events. The resulting DcaP classifications were visualized and annotated in iTOL (88). To infer a species phylogeny, a set of 120 universal bacterial marker genes (Bac120) was identified from each proteome using HMMER3 (hmmsearch, v3.3), following the methodology described in Parks *et al*. 2018 (89). The majority of species had only one significant hit for each of the 120 protein HMMs; if a species had more than one hit, the best-scoring hit for that marker was extracted.

Each of the 120 marker proteins was independently aligned to its respective HMM profile using hmmalign (HMMER v3.3). Alignments were trimmed to remove poorly aligned regions and concatenated into a single concatenated supermatrix for each species. To improve alignment quality, columns were filtered based on the following criteria: (i) highly gapped regions (>80% gaps) were removed; (ii) completely homogenous columns were excluded to reduce redundancy.

Maximum likelihood species trees were inferred from the filtered concatenated alignment using FastTree v2.1.11, with the LG model of amino acid evolution. The resulting unrooted trees were manually rooted and annotated in iTOL using *Brevundimonas subvibrioides* or *E. coli* as outgroups for the smaller and larger datasets.

### DcaP protein phylogeny

DcaP homologs were extracted from their respective proteomes by selecting sequences within the DcaP HOG at the root of all *Acinetobacter* species in the 255-species dataset in Orthofinder. To ensure consistent domain identification, sequences were aligned to the Pfam Hidden Markov Model (HMM) PF19577.4, corresponding to the DcaP OM protein family. Alignments were trimmed to remove poorly aligned residues prior to phylogenetic inference. A maximum likelihood phylogenetic tree was constructed using FastTree v2.1.11 with the LG model of amino acid evolution. The inferred clusters were identical to OrthoFinder’s DcaP subfamilies, except for a small cluster of DcaP proteins that OrthoFinder grouped with DcaP1, while FastTree grouped the same cluster with DcaP3.

To identify putative orthologs of DcaP proteins from the 22 species identified above and *A. baumannii* AB5075, proteomes from genomic assemblies were downloaded from the NCBI Datasets database. Two complementary approaches were employed to verify orthology. First, HMMER v3.4 was used to search each proteome with the Pfam profile HMM PF19577, corresponding to the DcaP protein family. Significant hits were retained as candidate DcaP homologs. To independently assess orthology and reconstruct gene relationships, OrthoFinder v2.5.5 was run on all 23 proteomes, which also produced a rooted gene tree for the DcaP family. The resulting DcaP sequences were realigned to the PF19577 HMM using hmmalign, and poorly aligned or unaligned amino acids were trimmed. A maximum likelihood phylogeny was then inferred using RAxML v8.2.12, under the LG amino acid substitution model.

Amino acid sequence alignment and percent identity were determined using the Clustal Omega program (1.2.4) through UniProt and the DcaP-family porin secondary structure was based on PDB entry 6EUS (28,90,91).

### Carbon source utilization screen using Biolog plates

Plates PM1 and PM2A were obtained from Biolog for carbon source utilization testing. A 110 µL aliquot of defined M9 minimal media lacking carbon was dispensed into every well and resuspended. Then, 100 µL was transferred from the Biolog plates into sterile flat-bottom 96-well plates. Overnight bacterial cultures were diluted 100-fold in sterile PBS before 1 µL of diluted culture was inoculated into every well. Plates were incubated at 37°C with shaking and growth was monitored by optical density at 600 nm (OD_600_) in an EPOCH2 or SynergyH1 BioTek plate reader (Winooski, VT). Background absorbance was subtracted before area under the curve calculations for each isolate and each compound were calculated and the difference between wildtype and ΔΔΔ*dcaP1-3* was analyzed. Compounds were included if any isolate of either wildtype or ΔΔΔ*dcaP1-3* reached an OD_600_ of 0.15 or higher.

### Growth curve analyses

M9 minimal medium without glucose was supplemented 0.1X Vishniac’s trace minerals (92) with the specified carbon source at a concentration to maintain the carbon equivalency of a 4-carbon compound at 16.5 mM. Cultures were inoculated with a single bacterial colony and grown in 3 mL LB at 37°C with shaking at 180 rpm for 8-16 hours. Growth curves were performed by diluting LB cultures 100-fold in sterile PBS and inoculating 1 µL diluted culture into 99 µL media in a 96-well flat bottom plate. Growth was monitored as described above.

### Murine bloodstream infection

Six-week-old female and male ND4 Swiss Webster mice were purchased from Envigo/Harlan Laboratory. Mice were boarded in a temperature-controlled environment with 14:10 h light:dark cycles and food and water were provided as needed. Mice were acclimated to the facility for 1 week prior to infection. Mice were anesthetized with isoflurane and inoculated retro-orbitally with approximately 3 × 10^8^ CFU in a 50 µL bacterial suspension of a 1:1 mixture of *A. baumannii* ATCC 17978VU wildtype and ΔΔΔ*dcaP1-3*::Kn mutant, ΔΔΔ*dcaP1-3*::Kn+*dcaP3*, or *ΔdcaP3*::Kn. For wildtype versus ΔΔΔ*dcaP1-3,* 5 of the mice were infected with wildtype ATCC 17978VU and 5 were infected with ATCC 17978 mTn*7*(Carb^R^); for the wildtype versus Δ*dcaP3*, wildtype was ATCC 17978 mTn*7*(Carb^R^). All mice infected with the complement strain were infected with ATCC 17978VU. The inoculum dose was determined by serial dilution and plating on selective agar media. Mice were euthanized at 24 h post infection by CO_2_ asphyxiation, and the organs were excised aseptically. Tissues were homogenized in PBS using a Bullet Blender (Next Advance, Troy, NY), and all samples were serially diluted and plated on LB and LB with carbenicillin or kanamycin selective agar plates for bacterial enumeration. All animal care protocols were approved by the University of Illinois Chicago Institutional Animal Care and Use Committee (IACUC; protocol number 23–119) in accordance with the Animal Care Policies of UIC, the Animal Welfare Act, the National Institutes of Health, and the American Veterinary Medical Association (AVMA). Animals were humanely euthanized consistent with the AVMA guidelines.

### Data reporting, statistical analysis, and figure preparation

Each measurement was taken from a distinct biological sample (e.g., bacterial culture from a single colony or an individual mouse). Data processing and statistical analyses were performed using Microsoft Excel 16.98 and GraphPad Prism 10.4.2 or 10.6.1. Statistical tests used are indicated in each figure legend. Figures were prepared in Adobe Illustrator 25.2.3 and 30.2.1.

## Author contributions

H.R.N. – project conceptualization, methodology, data collection, data analysis, and writing.

N.R.K. – data collection, data analysis, and writing. J.D.W – bioinformatics, data analysis, and writing. L.D.P. – project conceptualization, methodology, data analysis, writing, funding, and resources.

## ACKNOWLEDGMENTS

We thank the members of the Palmer laboratory for critical reading of the manuscript, Xiaomei Ren for construction of pLDP357, and funding from the National Institutes of Health R01AI189516 to L.D.P. and the Michael Reese Research and Education Foundation Pioneer in Research Award.

---

**Figure S1.**
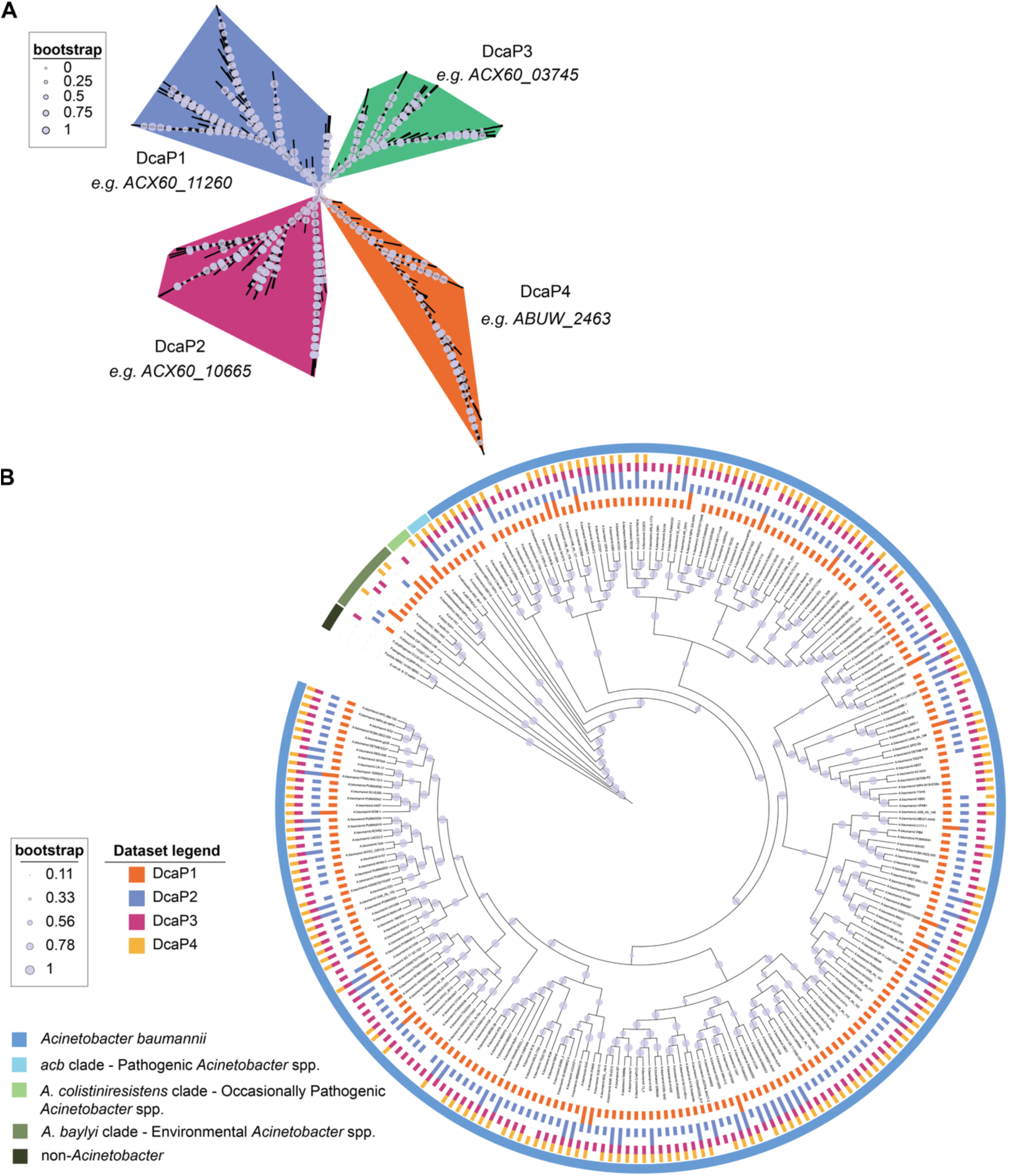
DcaP proteins are conserved across *Acinetobacter*. **(A)** *A. baumannii* DcaP proteins clustering with bootstrap values displayed. **(B)** Phylogenetic tree of *Acinetobacter* strains and clades depicting prevalence of individual DcaP proteins. Color coded boxes indicate the presence, absence, or duplication of individual DcaP proteins. Clades within *Acinetobacter* are indicated by the outermost color bar circling the tree. The accession numbers of the data depicted are in Table S4.

**Figure S2.**
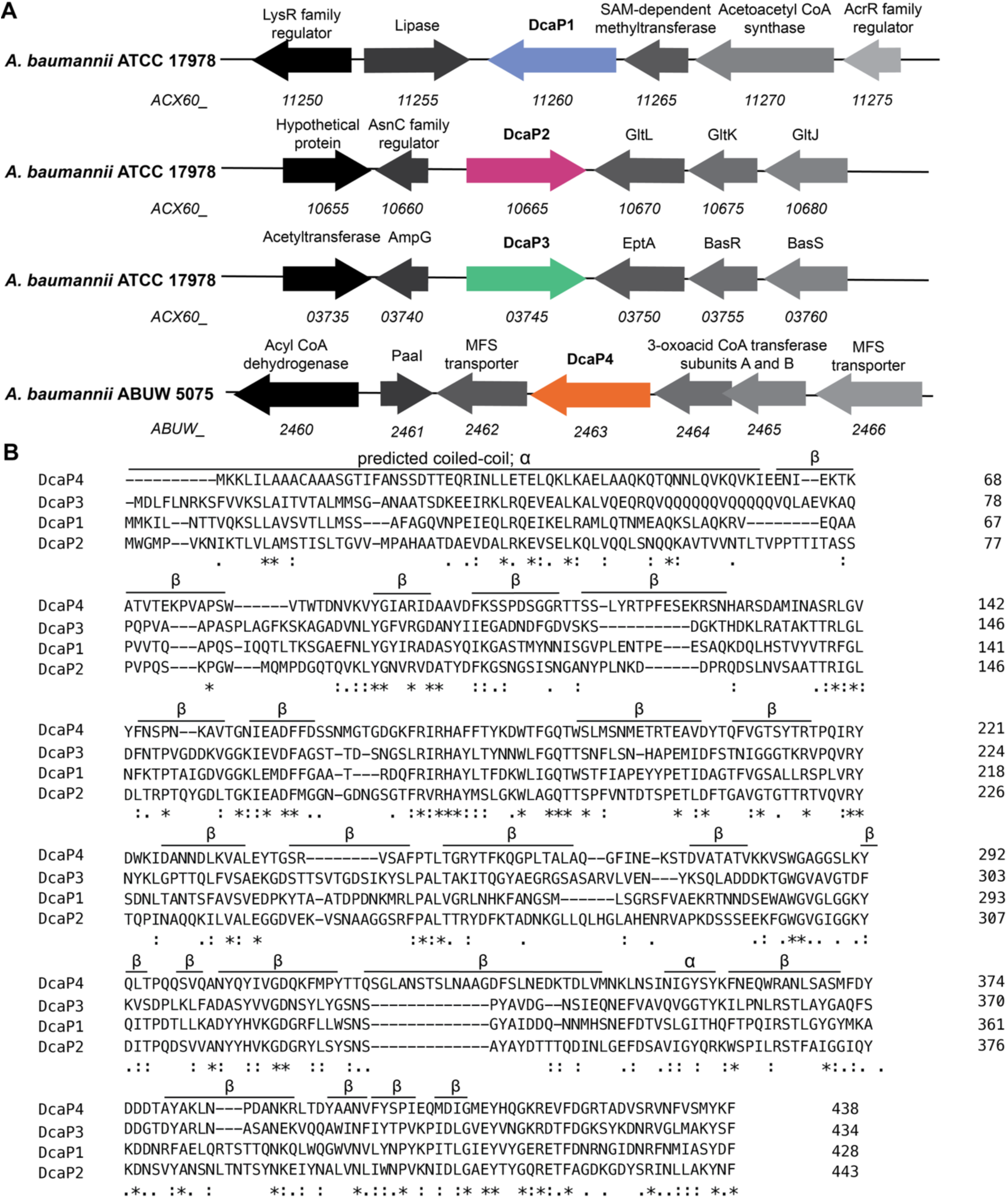
*dcaP* genomic neighborhoods and DcaP protein alignments. **(A)** The genomic loci of *dcaP1-4* are depicted. **(B)** CLUSTAL Protein alignment of DcaP proteins from ABUW 5075 (1). Secondary structure modeled after PDB entry 6EUS (2).

**Figure S3.**
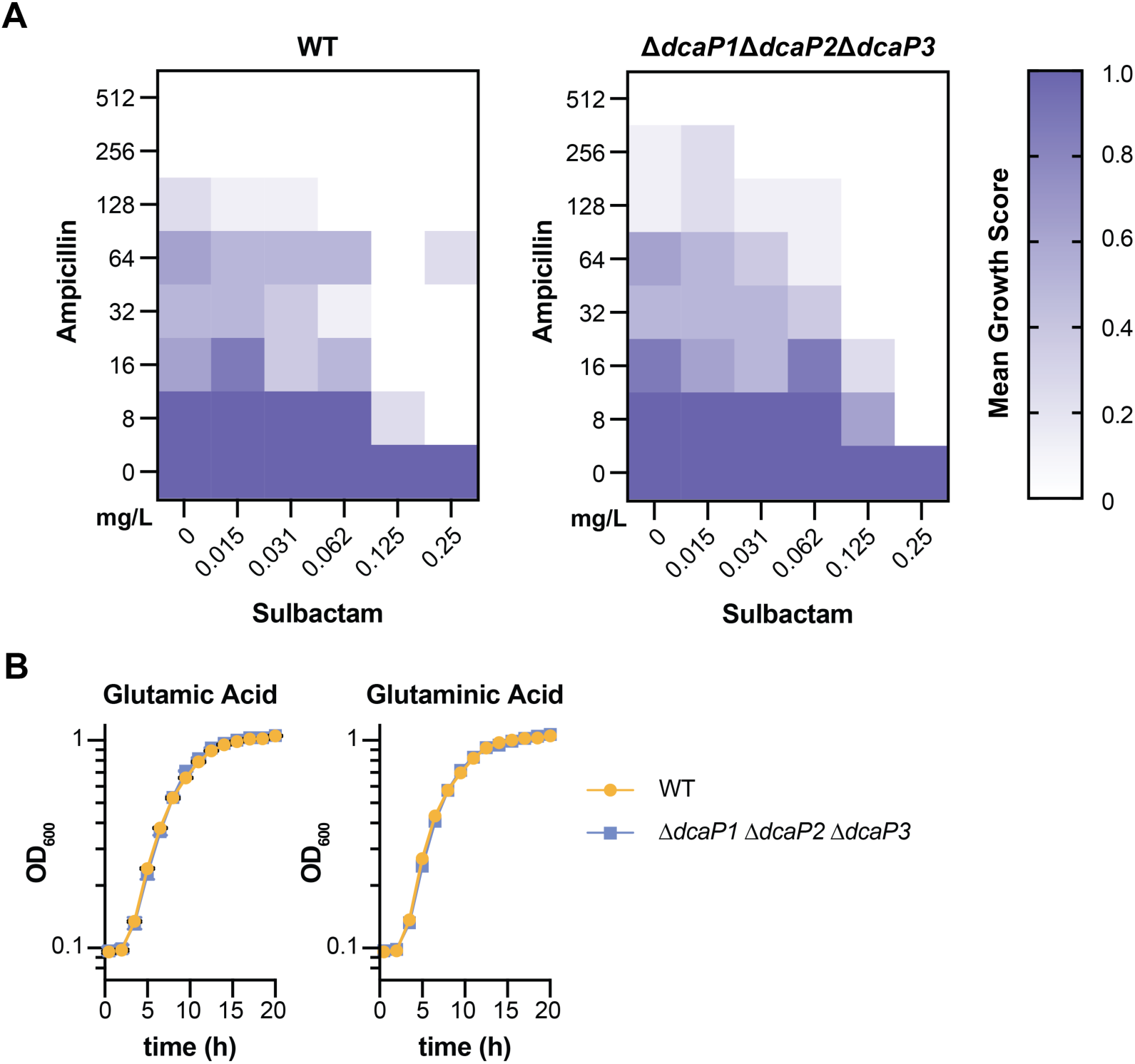
Checkerboard assay of ampicillin and sulbactam for wild-type and triple DcaP mutant and growth on glutamic acid and glutaminic acid. **(A)** A 96-well plate was filled with stocks of LB with ampicillin and sulbactam and serially diluted to obtain 100 µL of unique concentrations for each compound in each well before inoculation with 1 µL of indicated bacterial strain. Growth was scored as 1, strong turbidity; 0.5, mild turbidity; 0, little to no turbidity. Experiment was repeated 3 times with an n=2 for a total n=6. Means are shown. **(B)** Wildtype and the triple ΔΔΔ*dcaP1-3* mutant were grown in M9 media with the indicated compound as the sole carbon source. Growth was monitored by optical density at 600 nm (OD_600_). Data are mean +/- standard error of the mean (SEM). N=3 and data are representative of at least 2 independent experiments.

**Figure S4.**
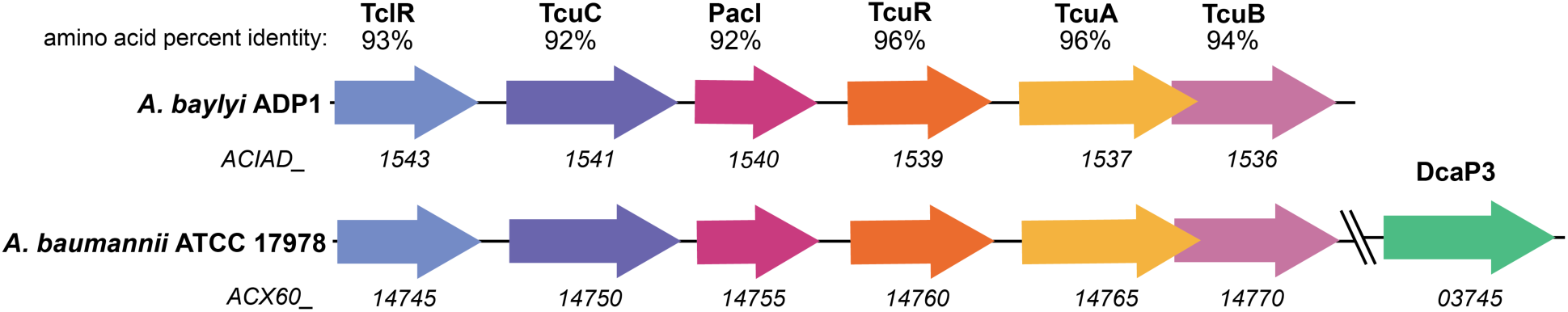
The genetic organization of the *tcu* metabolic locus in *Acinetobacter baylyi* ADP1 and *A. baumannii* ATCC 17978. Amino acid identity was determined by CLUSTAL Protein alignment using UniProt (1, 3).

**Table S1.** Strains.

| Strain | Source | Identifier |
| --- | --- | --- |
| *A. baumannii* ATCC 17978VU (WT) | ATCC | LP486 |
| *A. baumannii* ATCC 17978VU *att::mTn7*(empty); Carb<sup>R</sup> | Ren and Clark *et al.*, 2025 (4) | LP526 |
| *A. baumannii* ATCC 17978VU $\Delta dcaP1::kan$ (ACX60_11260) | This manuscript | LP1191 |
| *A. baumannii* ATCC 17978VU $\Delta dcaP2::kan$ (ACX60_10665) | This manuscript | LP1190 |
| *A. baumannii* ATCC 17978VU $\Delta dcaP3::kan$ (ACX60_03745) | This manuscript | LP1189 |
| *A. baumannii* ATCC 17978 $\Delta dcaP1\Delta dcaP2::kan$ | This manuscript | LP1253 |
| *A. baumannii* ATCC 17978VU $\Delta dcaP1 \Delta dcaP3 \Delta dcaP2::kan$ | This manuscript | LP1308 |
| *A. baumannii* ATCC 17978VU $\Delta dcaP3\Delta dcaP1::kan$ | This manuscript | LP1302 |
| *A. baumannii* ATCC 17978VU $\Delta dcaP3\Delta dcaP2::kan$ | This manuscript | LP1227 |
| *A. baumannii* ATCC 17978VU $\Delta dcaP1 \Delta dcaP3 \Delta dcaP2::kan$<br>*att::mTn7(dcaP3p-dcaP3)* | This manuscript | LP1310 |
| *A. baumannii* ATCC 17978VU $\Delta dcaP1 \Delta dcaP3 \Delta dcaP2::kan$<br>*att::mTn7(dcaP3p-dcaP4(ABUW_2463))* | This manuscript | LP1314 |
| *A. baumannii* AB5075 | Colin Manoil (University of Washington) (5, 6) | LP345 |
| *A. baumannii* AB5075 $\Delta dcaP3::hyg$ (ABUW_RS04045) | This manuscript | LP1315 |
| *A. baumannii* ATCC 17978VU $\Delta dcaP1 \Delta dcaP3 \Delta dcaP2::kan$<br>*att::mTn7(dcaP3p-dcaP1)* | This manuscript | LP1368 |
| *A. baumannii* ATCC 17978VU $\Delta dcaP1 \Delta dcaP3 \Delta dcaP2::kan$<br>*att::mTn7(dcaP3p-dcaP1)* | This manuscript | LP1369 |

**Table S2.** Plasmids.

| Use or Phenotype | Source | Name | Identifier |
| --- | --- | --- | --- |
| Kan<sup>R</sup>; mobilization helper plasmid | Figurski and Helinski, 1979 (7) | pRK2013 |  |
| Carb<sup>R</sup>; plasmid encoding mTn7 transposition pathway | Choi *et al.*, 2005 (8) | pTNS2 |  |
| Carb<sup>R</sup>; allelic exchange vector with sucrose sensitivity | Hoang *et al.*, 1998 (9) | pFLP2 |  |
| Contains an FRT-flanked kanamycin resistance gene | Datsenko and Wanner, 2000 (10) | pKD4 |  |
| Contains a mTn7 with carbenicillin resistance gene | Carruthers *et al.*, 2013 (11) and Alexeyev, 1999 (12) | pKNOCK |  |
| Carb<sup>R</sup>; construct to complement *dcaP3* | This manuscript | pKNOCK-*dcaP3* | pLDP343 |
| Carb<sup>R</sup>; express the AB5075 *DcaP4* in ATCC 17978 | This manuscript | pKNOCK-*dcaP4* | pLDP359 |
| Carb<sup>R</sup> Kan<sup>R</sup>; deletion construct for *DcaP1* | This manuscript | pFLP2-*dcaP1* | pFLP2-11260 |
| Carb<sup>R</sup> Kan<sup>R</sup>; deletion construct for *DcaP2* | This manuscript | pFLP2-*dcaP2* | pFLP2-10665 |
| Carb<sup>R</sup> Kan<sup>R</sup>; deletion construct for *DcaP3* | This manuscript | pFLP2-*dcaP3* | pFLP2-03745 |
| pKD4 with hygromycin<sup>R</sup> replacing kanamycin<sup>R</sup> cassette | This manuscript | pKD4-Hyg | pLDP357 |
| Carb<sup>R</sup> Hyg<sup>R</sup>; deletion construct for *DcaP3* in AB5075 | This manuscript | pFLP2-5075*dcaP3* | pLDP360 |
| Carb<sup>R</sup>; construct to express *dcaP1* | This manuscript | pKNOCK-*dcaP1* | pLDP401 |
| Carb<sup>R</sup>; construct to express *dcaP2* | This manuscript | pKNOCK-*dcaP2* | pLDP402 |

**Table S3.**
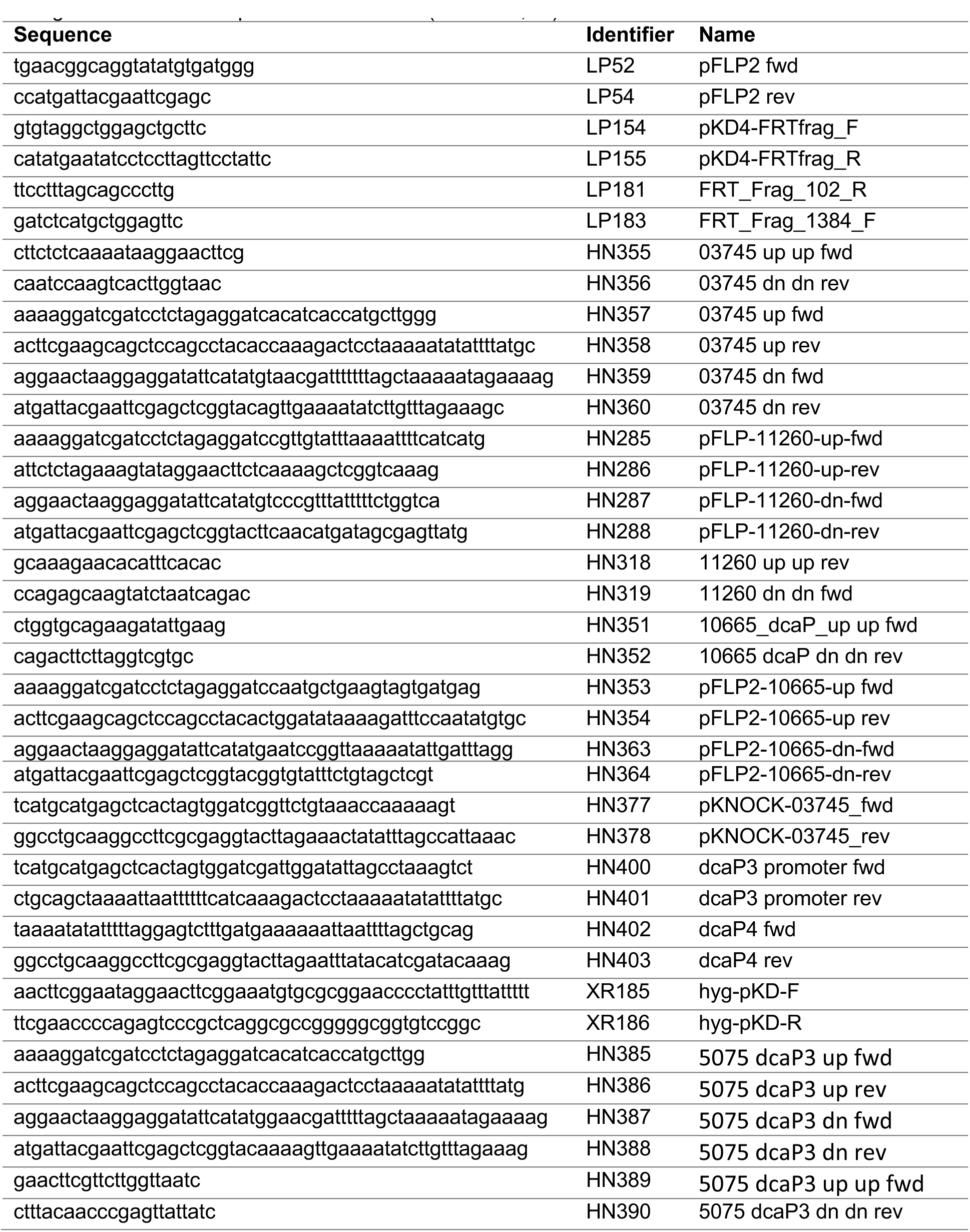
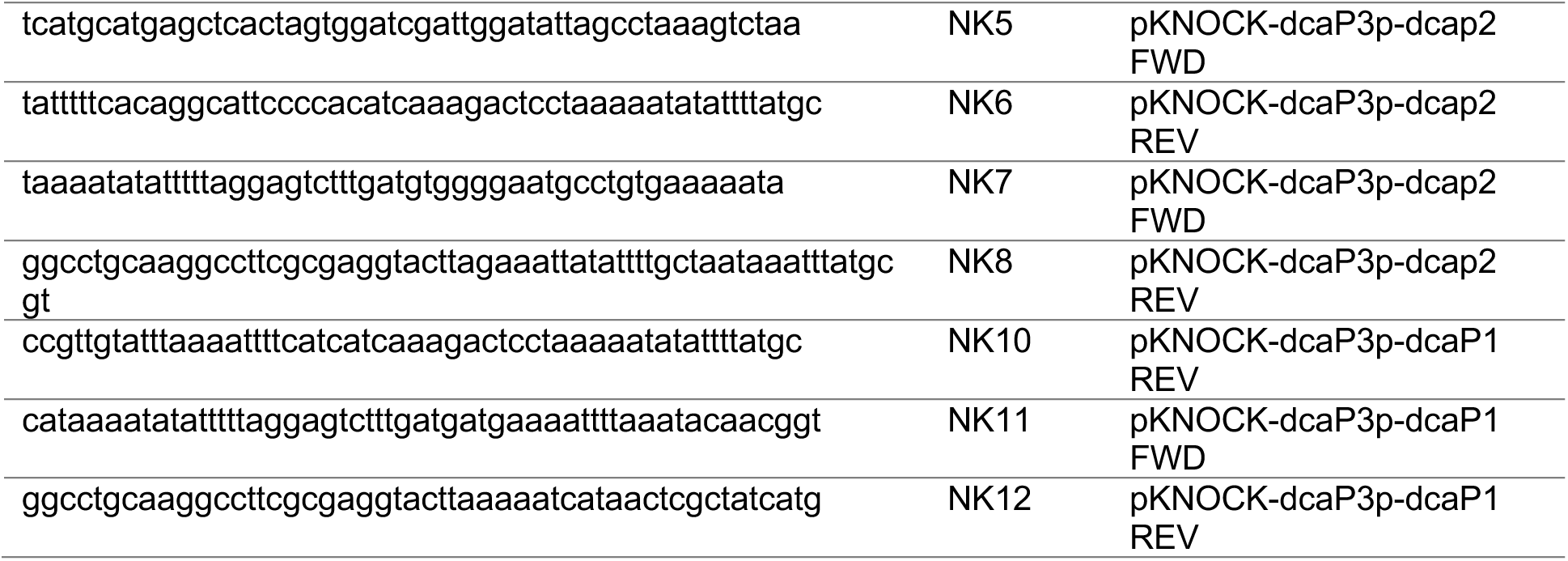
Oligonucleotides. All oligonucleotides were purchased from IDT (Coralville, IA)

## Supplementary Methods

### Checkerboard assay

In a 96-well plate, each well was filled with 100 µl LB. Stock solutions of ampicillin (100 mg/mL) and sulbactam (30 mg/mL) were diluted into LB for a final concentration of 1024 µg/mL and 32 µg/mL, respectively. An additional solution of 2048 µg/mL ampicillin was prepared. 100 µl aliquots of 1024 µg/mL ampicillin were placed in row A, wells 1-11, and 2024 µg/mL ampicillin in well 12. 2-fold serial dilutions were performed beginning in row A and ending in row H, discarding the remaining 100 µl. 100 µl aliquots of 32 µg/mL sulbactam were placed in every well in column 12, and 2-fold serial dilutions were performed beginning in column 12 and ending in column 2, discarding the remaining 100 µl. The maximum concentration for each antibiotic was therefore 512 µg/mL ampicillin and 16 µg/mL sulbactam. Then, 1 µl overnight culture of *A. baumannii* ATCC 17978 wild-type or the ΔΔΔ*dcaP* mutant was inoculated into each well before incubation at 37°C with shaking at 180 rpm for 8-16 hours. The following day, plates were scored for growth or no growth phenotypes. Wells with strong optical turbidity were assigned a value of ‘1’ and wells with mild turbidity or mild turbidity with clumped cells were assigned a value of ‘0.5’. Wells with no visual growth or fully clumped cells were assigned a value of ‘0.’

